# Phage phenotyping by measuring plaque expansion dynamics

**DOI:** 10.64898/2026.10.07.757331

**Authors:** Jonathan Costa, Baltus A van der Steen, Yuval Mulla

## Abstract

Bacteriophages exhibit staggering genomic diversity, yet phenotypic characterization remains bottlenecked by assays that generally yield simple binary infection outcomes. Quantitative traits can be extracted from standard liquid cultures, but many phages fail to produce a measurable effect in these settings, necessitating an improved quantitative approach on semi-solid media. While end-point plaque size is commonly used for quantitative phenotyping, it only provides a static snapshot that fails to disentangle phage performance across changing bacterial growth phases. Here, we report a cost-effective, scalable method to quantify the bacterial killing rates of diverse phages by continuously tracking plaque development using a consumer-grade flatbed scanner and custom open-source computer vision software. By fitting a minimal, phenomenological model to the diverse BASEL collection of *Escherichia coli* phages, we accurately captured plaque expansion dynamics across all bacterial growth phases. We discovered that plaque expansion parameters vary significantly across individual phages and taxonomic families, with the initial expansion rate demonstrating an inverse correlation with viral genome size. Notably, these plaque expansion dynamics did not correlate with bacterial collapse times in liquid culture, indicating that they represent distinct physiological and kinetic processes. Finally, we demonstrated that phage taxonomic family can be predicted with 72.1% accuracy solely from dynamic plaque parameters using a random forest classifier, whereas conventional endpoint plaque sizes held no predictive power. This time-lapse phenotyping approach yields robust kinetic data superior to end-point measurements, establishing a scalable platform to accelerate phage discovery and genotype-phenotype mapping.

## Introduction

Bacterium-infecting viruses (phages) exhibit staggering diversity^1^. Metagenomic sequencing has identified over 150,000 distinct phage species^2^, a figure nearly two orders of magnitude greater than the number of bacterial species discovered via the same approach^3^. Phenotypically, this diversity is equally profound. Phages display high specificity for their bacterial hosts, with host ranges varying between phages due to a range of factors such as divergent receptor specificities and sensitivities to host immune systems^4^. Furthermore, even different phages targeting the same host exhibit drastic variations in their rate of infection^5^.

While advances in metagenomic sequencing have made it routine to capture genomic diversity at scale, experimental methods for quantitatively phenotyping phages remain bottlenecked. Most phenotypic characterizations are restricted to a binary classification of whether a phage can or cannot infect a specific bacterium^6–10^. Differences in the rates of infection are entirely missed by these binary approaches. Quantifying this bacterial killing rate is not only of fundamental scientific interest but is also clinically relevant, as it serves as a critical determinant of therapeutic success in phage therapy^11–13^.

Current methodologies of quantitatively inferring phage activity mostly rely on monitoring phage predation in a bacterial population growing in liquid culture^5,14,15^. However, many phages capable of successfully replicating on and killing a host fail to produce a measurable effect in liquid suspensions^16,17^. As a result, the killing rates of many diverse phages cannot be accurately quantified in liquid media.

The gold standard for evaluating phage infectivity is the plaque assay. In this setup, a dense bacterial lawn is established within a semi-solid agar layer, and a phage population, originating from a single viral particle, expands radially as it lyses and replicates on the host^13^. While scoring a binary infection/non-infection outcome is straightforward^6–10^, quantifying the actual bacterial killing rate from plaque assays is more problematic.

Absolute phage yield can be quantified by counting plaques, but viral yield does not correlate with the bacterial killing rate across different phages on the same host^5^. Plaque counting is also used to determine the efficiency-of-plating (EOP), which compares the number of infective particles between a reference host and a target host^18^. While EOP allows researchers to compare how frequently an infection succeeds across different hosts, it does not capture differences in infection *kinetics*.

Alternatively, researchers have long used endpoint plaque size and morphology to compare different phages or to assess the effects of stressors like antibiotics^19–21^. While plaque size is easy to measure, it offers only a static, endpoint snapshot of a highly dynamic process. This poses a distinct limitation because a bacterial population undergoes sequential growth phases, spanning lag, exponential growth, and stationary phase, while phages display unique kinetics within each phase^22^. Consequently, endpoint plaque size acts as a composite metric that fails to disentangle phage performance across these changing bacterial growth states^23^.

Capturing bacterial killing across different growth phases requires tracking plaque dynamics over time rather than relying on final sizes alone. Significant theoretical work has modeled the plaque dynamics of the T7 phage after host bacteria have entered stationary phase^24–27^. This late phase is relatively easy to track experimentally due to the high contrast between lysed and unlysed regions on the bacterial lawn, alongside the slow, steady-state dynamics that unfold over several days^28–30^. However, most phages exhibit little to no replication dynamics during this late phase, replicating exclusively on actively growing bacteria^31^. Despite sustained theoretical interest^32,33^ and the need to phenotype phages that are inactive in liquid culture, we still lack dynamic plaque expansion data for diverse phages on actively growing hosts^34,35^.

Here, we report a cost-effective, scalable approach to quantify the bacterial killing rates of diverse phages on hosts growing in semi-solid media. By spotting phages onto a bacterial lawn, we track plaque development using a consumer-grade flatbed scanner and custom, open-source image analysis software. We demonstrate that a minimal phenomenological model fully captures plaque dynamics across all bacterial growth phases. Applying this model to the BASEL collection, a diverse reference set of phages infecting *Escherichia coli* MG1655^31^, we show that the plaque expansion parameters vary significantly across both individual phages and taxonomic families. Finally, we demonstrate that phage family identity can be accurately predicted solely from plaque dynamics, whereas conventional endpoint plaque size measurements hold no predictive power.

## Materials and methods

### Bacterial strains and phages

The bacterial strain used as a host was *Escherichia coli* K-12 MG1655. All phages used come from the BASEL phage collection^31^ and are numbered exactly as they appear in the collection.

### Media and agar

The LB (Lysogeny Broth) media used in all experiments was prepared from BD Difco™ LB Broth Lennox (BD Biosciences, 240230) powder. The agar used to prepare the plates and phage dilutions was prepared from BD Difco™ LB Agar Lennox (BD Biosciences, 240110).

### Plaque creation

Double-layer LBA within single-well plates (thermoscientific, OmniTray 242811) was created using 10 mL 1.5% agar for the bottom layer and 5 mL 0.5% agar for the top layer. Before pouring the top layer, an _E. coli_ overnight culture (incubated for 20 h at 30 °C in LB media in a shaker flask at 200 rpm) at an extrapolated OD_600_ of 5 was added at a dilution of 1:30. These plates were then incubated at 30 °C for 2.5 h.

After this first incubation, 3 µL of phage solution was spotted onto the bacterial lawn. To capture plaques from single phage origin, the stock solution was diluted to a concentration between 1 - 25 PFU (plaque forming units) per 3 µL. The exact concentration was chosen per phage depending on the size of its plaques. To maximize data yield, the highest possible PFU concentration that still prevented individual plaques from merging was preferred. Additionally, 2 µL Streptomycin (15 mg mL^−1^) was added to a phage-free area of the plate to provide a background correction reference. The spotted solution was left to dry for 5 min. Afterwards, the plate was sealed with Parafilm to prevent desiccation and placed on the scanner inside the incubator at 30 °C for time-lapse scanning.

To prevent condensation, the lids of the single-well plates were coated with a solution of 20% (v/v) Ethanol and 0.05% (v/v) Triton X-207^36^. The inner lid surfaces were covered with 3 mL of this solution and incubated for 30 s. Then the excess liquid was shaken off, and the lid was left to air-dry completely.

### Time Series Scanning

Scans were performed using an Epson Perfection V600 Photo scanner (B11B198032) controlled via custom software built on top of the standardized TWAIN interface^37^. To communicate with the TWAIN interface, the UniteLabs connector development toolkit^38^ was used to create a software package following the SiLA 2 standard specifications^39,40^. SiLA 2 is a standard built for laboratory device integration, which was chosen here to make the image capture process accessible for use in larger automated work cells. This software has been open-sourced^41^.

For this work, scans were captured as 24-bit RGB images at a resolution of 1500 dpi with the scanner set to transmission mode. While this setting corresponds to a theoretical pixel size of 0.016933 mm, the plaques were imaged slightly above the scanner’s focal plane. This resulted in optical blur, reducing the effective resolution well below this theoretical 0.01 mm limit. Because of this blurring effect, we encountered difficulties accurately imaging the very small plaques present in the phage library, and increasing the scanner’s hardware resolution further did not resolve this issue. Although this optical limitation does not negatively affect the internal consistency of the results of the resolvable plaques presented here, it may limit direct quantitative comparability with data obtained using different optical setups. However, we postulate that relative results should remain comparable across setups.

Scanning at a frequency of one image every 10 min for 16 h produced 41 GB of raw pixel data per run. This data, along with experiment metadata, was compiled into an xarray dataset^42^, then compressed and stored using the zarr format^43^. This format was chosen because it allows for the efficient storage and retrieval of large multi-dimensional arrays without requiring the entire dataset to be loaded into memory at once.

### Image Pre-processing

Raw pixel data was processed using a custom-built Python package, which is open-sourced^41^. The package provides a general set of tools for processing, analysing, and visualizing plaque image data, establishing an extensible framework for phage plaque analysis. Its structure is strongly inspired by the data analysis and manipulation library ‘pandas’ to provide an interface familiar to scientists accustomed to data processing in Python. All package dependencies are open source and can be found in the ‘pyproject.toml’ file. Version control is managed via ‘git’ and compatibility is ensured using ‘uv’^44^ for dependency and virtual environment management. The exact package versions used, and a quick installation guide are available in the Appendix.

The resulting images were manually annotated. This means selecting the perimeters of the plates, single plaques, and correction areas for both undisturbed bacteria and background. This spatial “chunk” selection is done by using a custom-built user interface running in a marimo notebook. For the purposes of this work, the raw pixel intensity *I*_0_ was first inverted so that lower intensity is equivalent to the absence of bacteria (Eq. 1), and then scaled against the inverted intensities of both the bacteria *I*_bacteria_ and the background *I*_background_ to increase the contrast (Eq. 2). For this scaling, the closest available correction chunks were automatically selected by the software. Finally, the pixel values were normalized to a range of 0 to 1 (Eq. 3). While this normalization is not strictly necessary and does not affect the final mathematical results, it facilitates the observation and comparison of intermediate processing steps across different experiments. This process is also visualized in Fig. 1a.

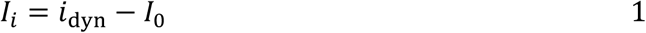

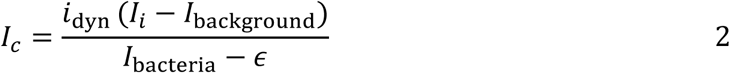

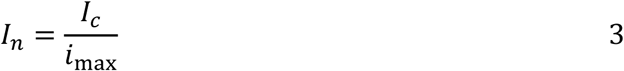

where *i*_dyn_ is the maximum intensity of the scanner’s dynamic range, *ε* is a very small constant to avoid division by zero and *i*_max_ is the maximum intensity within the current frame.

**Figure 1.**
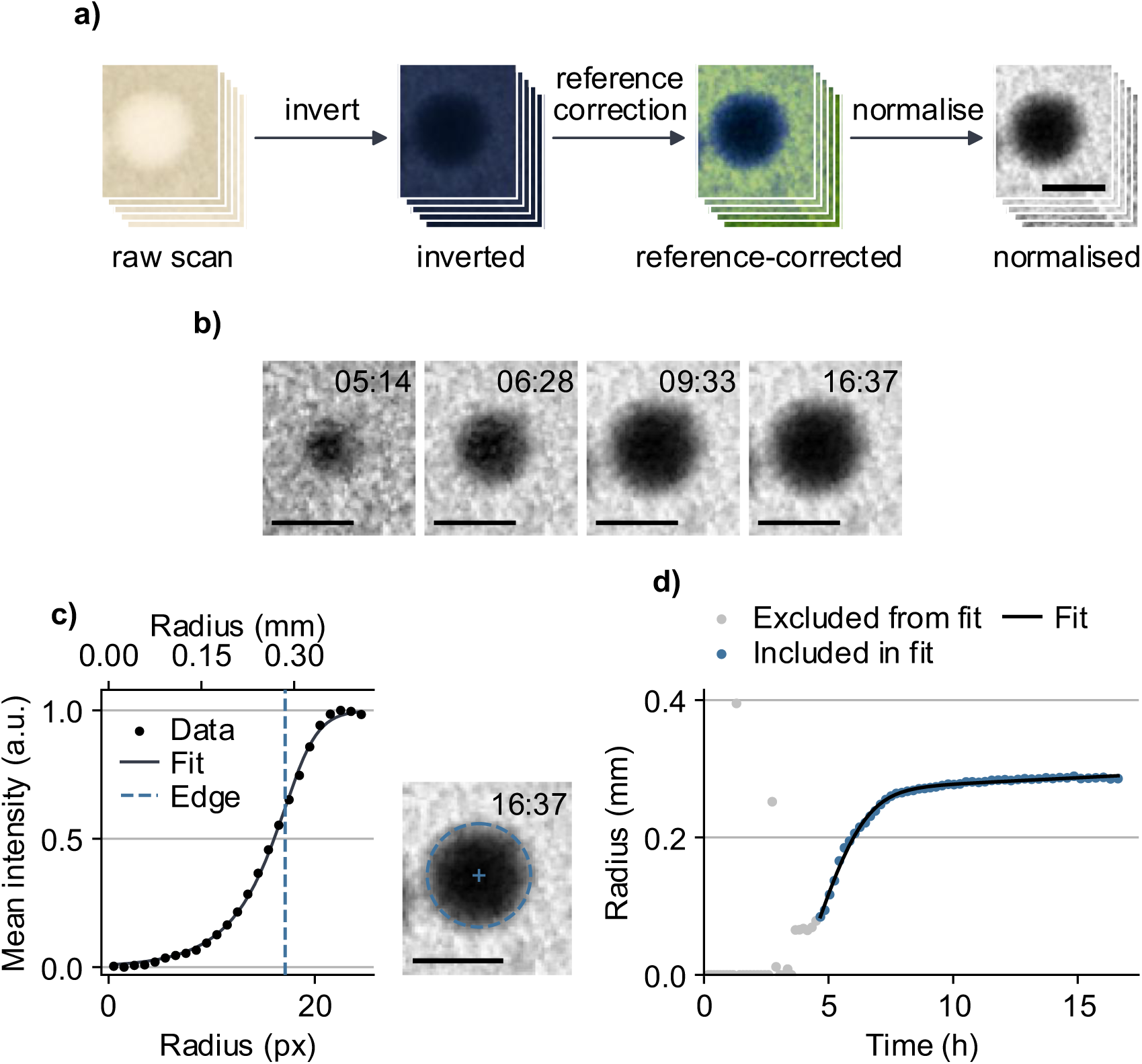
Computer vision tracks plaque radius over time. Expansion dynamics are shown for a plaque cleared by phage 31. **a)** Images are preprocessed to enhance visual contrast (methods). **b)** Four sample images from different timepoints out of all captured images of the plaque showing how contrast and size of the plaque increase over time. The length of the scale bar is 1 mm. The incubation time since the application of the phage is shown as hh:mm. **c)** Left: A sigmoidal fit (solid black line) is applied to the radially integrated plaque intensity profile measured at 1c:37 (dots) to find the plaque edge (blue dashed line). Right: visual inspection shows that the algorithm accurately detects the plaque edge (highlighted in green). **d)** Edge detection is applied over the entire time-series. The expansion dynamics are fitted with a phenomenological model (methods) to extract the rate of plaque expansion. Because the contrast between plaque and non-plaque is very weak in the beginning due to limited bacterial growth, the starting data points are excluded based on predetermined criteria (methods).

Uppercase *I* variables refer to a matrix of pixel values.

### Expansion Measurement and Model

After pre-processing, the time series of each plaque chunk was analysed to detect circular structures using Canny edge detection and the Circle Hough Transform from scikit-image^45^. The resulting circle parameters were used to calculate the plaque’s centre. These centre coordinates were then stabilized across all timepoints by assigning a custom confidence score to the estimated circle. This score was determined by verifying whether the area outside the estimated circle was significantly brighter than the inside. Using these stabilized centres, radial integration was performed at each timepoint, generating a 3D array of intensity as a function of radius and time. On each time slice of this array, a Richards curve was fitted (see Eq. 4), adapted with a lower boundary of 0 and upper boundary of 1, which after normalization (Eq. 3) are our lowest and highest expected intensities, respectively. This is a generalized sigmoidal function that was found to best fit the radial intensity profiles observed.

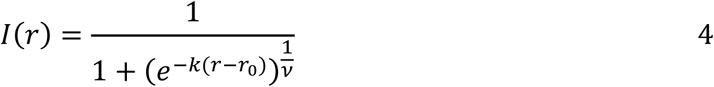

where *I*(*r*) is the intensity at radius *r*, *k* steepness, *r*_0_ marks the inflection point, and *ν* a constant determining how asymmetric the curve is.

From each timepoint the fitted *r*_0_ value (radius at the inflection point) was extracted to establish a radius-over-time profile representing plaque expansion. The radius was specifically derived from this radial integration and Richard’s curve fit, rather than relying directly on the initial Hough circle radii, because this yielded significantly less noise in the final expansion data.

These expansion curves were subsequently fitted using an empirical model consisting of two linear expansion phases connected via a logarithmic transition (see Eq. 5). This is not a biologically derived mechanistic model, but rather a phenomenological one based on qualitative observations of the expansion curves (Fig. 1d).

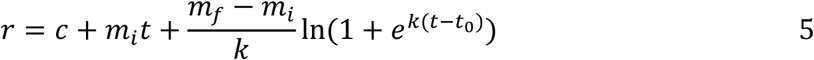

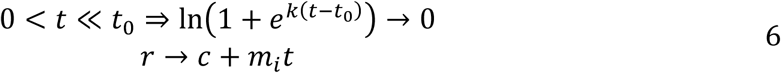

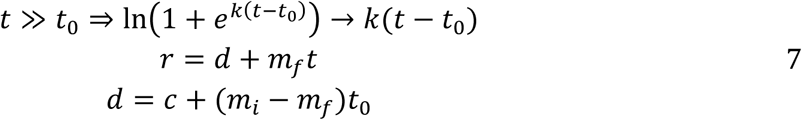

The following limits have been imposed on the curve fitting algorithm when estimating the parameters for the expansion model:

- *c* in [-∞, max(*r*_measured_): the upper limit is set to the max of the measured radius just to reduce compute as the y-axis intercept cannot exceed the maximal measured radius
- *m_i_* in 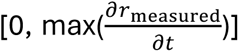 the lower bound of 0 was chosen to ensure expansion and the upper bounds similar to *c* are set at the maximal reachable within our dataset to reduce compute
- *m_f_* in 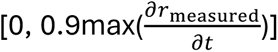 the upper bound is set to be smaller than the initial expansion rate to enforce a small change in expansion rate. In a case where the final rate is the same as the initial rate, we cannot distinguish two phases. No plaque has been observed that increases in expansion rate in the second phase.
- *t*_0_ in [5,11]: This means that the transition is restricted to fall somewhere between 5 h and 11 h. All observed phages have transitioned within this window.
- *k* in [1,10]: This parameter controls the speed of the transition. The chosen range allows for variability across plaques but restricts unnaturally fast and slow transitions.

Applying the correction, the radius extraction and the expansion model on each plaque in our database yielded a collection of initial and final expansion rates, as well as a theoretical expansion start.

### Quality filter

Within the described pipeline, there are four mathematical quality filters where some of the data are excluded automatically.

1. Circle Hough Transform: The quality of the circle *q*_circle_ estimated by the Circle Hough Transform is calculated by subtracting the mean intensity of all the pixels within the circle from the mean intensity of all the pixels outside the circle (Eq. 8). At this point all the pixel intensities have been normalized between 0 and 1, which makes 0 ≤ *q*_circle_ ≤ 1, where 1 is a circle with high contrast between inside and outside. The cutoff point in this work was set at 0.5. This means every frame where the *q*_circle_ < 0.5 is ignored. If none of the frames of a selected plaque shows *q*_circle_ ≥ 0.5, the plaque is removed from the database.

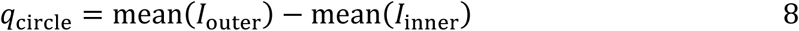
2. Sigmoidal fit: For each sigmoidal fit of a frame’s radial intensity profile, an *R*^2^ value is calculated from the resulting curve parameters. This *R*^2^ is used as a filter for which datapoints to consider for the expansion model fit. Inflection points that were extracted from a sigmoidal with *R*^2^ < 0.95 are excluded. This filter serves to exclude early measurements while the contrast between bacteria and plaque is not high enough to estimate the border and late measurements in case plaques start overlapping.
3. Model fit: If no data is available (not measured or excluded due to bad sigmoidal fit) after the transition window (5 to 11 h) to fit a final expansion phase, only an initial phase is fitted (as described in Eq. 6). If not enough data is available to fit the initial phase, the plaque is removed from the database.
4. Model fit quality: After fitting all the parameters of the expansion model (Eq. 5), three *R*^2^ values are calculated to report the quality of the fit. One 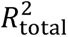 is calculated for the full model fit. Models that report a 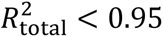 are removed from the database. As shown in Eq. 6 and Eq. 7, the model can be separated into two linear phases. For each of those phases, an individual 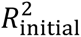 and 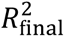 is calculated. As these are partial models, their filter is softer, with 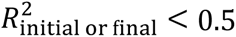 leading to removal of the individual phases from the database, respectively.

### Bacteria and phage propagation

*Escherichia coli* MG1655 carrying a luciferase-based bioluminescence reporter encoded on the kanamycin-resistant plasmid pCS-λ was used for all experiments. Pre-cultures were grown overnight at 30°C in high-salt lysogeny broth (LB) with shaking at 150 rpm for liquid cultures or on 1.5% LB agar (LBA) plates for solid cultures. Phages used in this study are part of the BASEL phage collection and propagated on *E. coli* in LB at 30°C with shaking (150 rpm) overnight. Phage lysates were purified by chloroform treatment (1:75, v/v), followed by centrifugation at 2800 × g.

### Single-phage growth curve

To characterize phage induced collapse times for the BASEL collection, *E.coli* was grown to the mid-exponential phase, diluted to an OD_600_ of 0.05 (1.69 × 10^7^ CFU/ml), and infected with 1000x diluted BASEL phage (1:200, v/v). Cultures were incubated in a 96-well plate format (200 μl/well) with double orbital shaking at 30°C for 48 h. Collapse time was determined by measuring OD_600_ and bioluminescence (BioTek Synergy HTX Multi Mode Plate Reader) every 10 min. Collapse times for both OD_600_ and bioluminescence curves were obtained by using find_peaks SciPy 1.1.13.1) with a smoothing width of three data points.

## Results

To capture phage dynamics on actively growing bacteria, we first incubate a lawn of *E. coli* before spotting a phage lysate on top of the agar. We spot a dilution series of the phage lysate to ensure plaques with a single-phage origin. Next, we incubate the plate at 30 degrees and image every 10 min for 16 hours to observe the formation of the phage plaques at the locations of spotting. Initially, the plate is homogeneous as the bacteria are to significantly scatter light. As the bacterial population grows denser, phage plaques become increasingly visible and expand (**Figure 1**a,b). We identify the center of each plaque (Methods) on the final image as it has the best contrast between plaque and non-plaque areas. We radially integrate the intensity from this center for each timestep to identify the plaque edge and calculate the plaque radius over time (**Figure 1**c). Across all traceable plaques we find reproducible dynamics: the plaque is poorly visible and hardly grows within the first few hours, after which it linearly expands with a rate that drops as the bacterial growth declines (**Figure 1**d, **Supplementary Figure *1***).

We next aim at capturing the dynamics of plaque expansion in a minimal set of phenomenological parameters. To fully capture the dynamics, we find 5 parameters optimal: we fit the initial lag phase with a delay time (1), and the resultant growth with an initial (2) and final expansion rate (3). Lastly, we include a transition time (4) and transition width (5) between the initial and final expansion (methods Eq. 5). We find that these five parameters accurately capture the full expansion dynamics (**Figure 1**d), two of which (delay time and initial rate) characterize the plaque expansion dynamics on bacteria growing at steady state.

To test the broader applicability of our plaque tracking and fitting algorithm, we applied it to the entire BASEL collection of 69 different lytic E. coli phages (**Figure 2**a). Out of the 69 phages, the titer of four phages was below any detectable level, five phages showed abnormal plaque morphology (large variety of plaque size and/or no clear borders), and thirteen had plaques which remained too small to track (**Supplementary Figure 2**). All remaining 47 phage plaques showed dynamics consistent with afore-mentioned dynamics.

**Figure 2.**
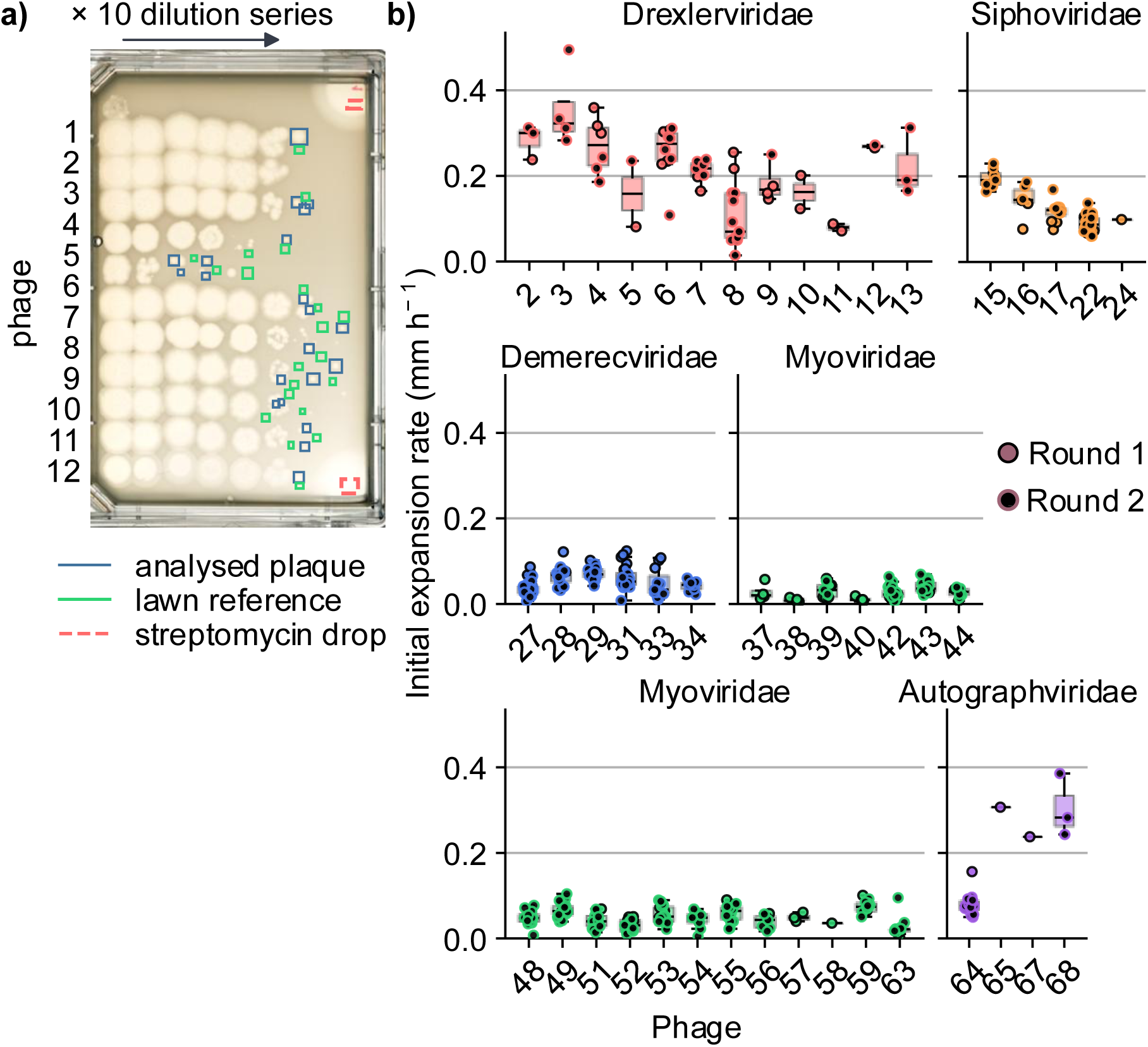
High-throughput spotting reveals diversity of plaque expansion dynamics across phages. **a)** Different phages (rows) were serially diluted (columns) and spotted on an agar plate. Plaques were selected manually for tracking expansion dynamics. The individual plaques are highlighted in different colors according to their phage of origin. Nearby non-plaque lawn areas were selected for reference intensity. **b)** Box plot showing the initial expansion rate for each phage and the phages are colored according to family. Data were collected over 2 rounds of experiments.

**Figure 3.**
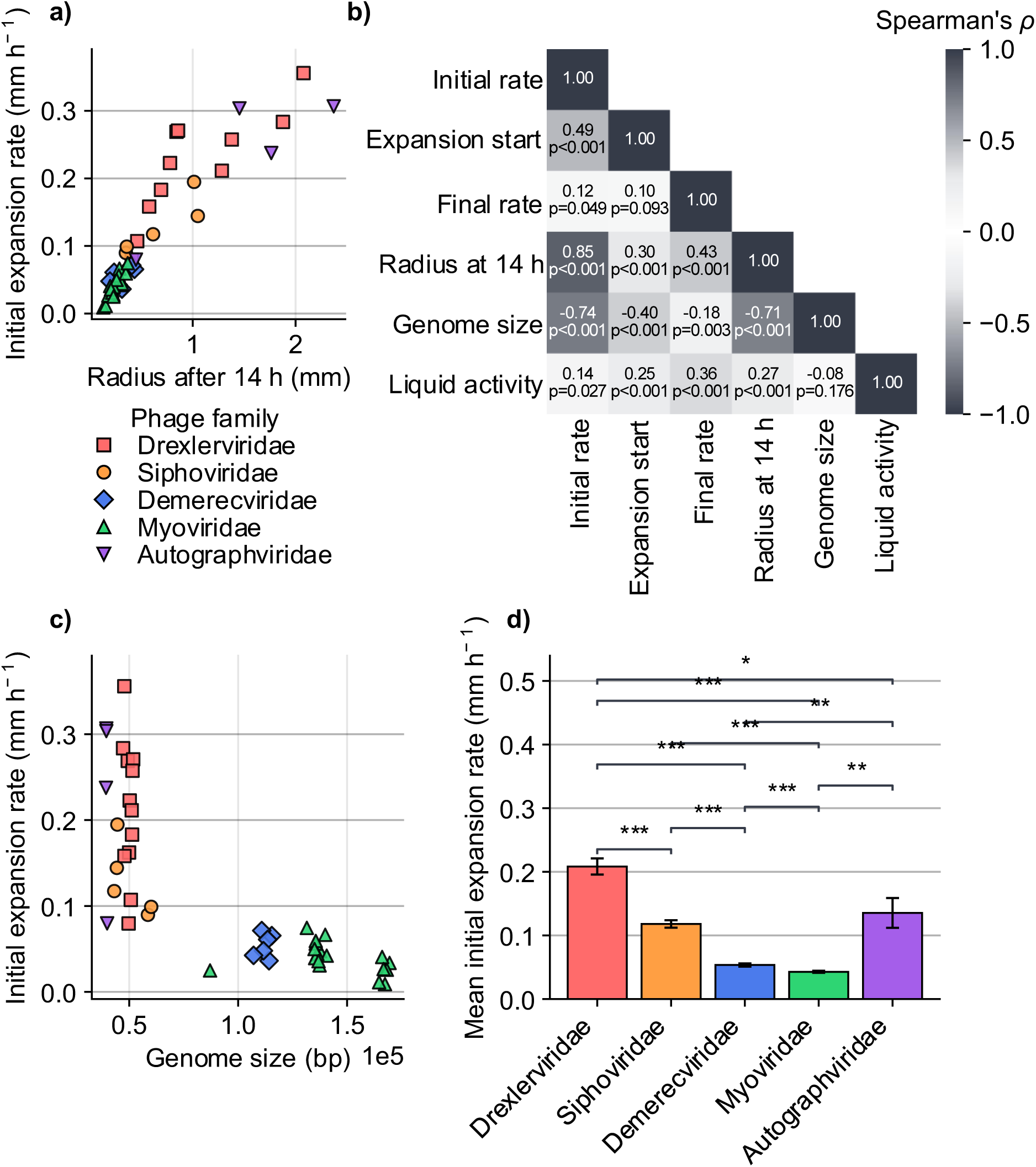
Determinants of plaque expansion dynamics. **a)** The initial plaque expansion rate correlates with the final genome size both across and within phage families. **b)** All vs. all spearman rank correlation across the BASEL phages, correlating the plaque expansion parameters, phage genome size and activity in liquid as defined by the collapse time. **c)** The initial expansion rate decays as a function of genome size, however the correlation between genome size and phage family (colors) is a confounding factor**. d)** The initial expansion rate (mean ± standard error) differs significantly among most families. For all panels, calculated p-values were adjusted for multiple comparisons using the Benjamini-Hochberg procedure (to avoid Type I errors). Asterisks denote adjusted significance levels (∗ p < 0.05, ∗∗ p < 0.01, ∗∗∗ p < 0.001).

To study phenotypic diversity, we extracted the plaque expansion dynamics across different phages. We find that the initial expansion rate is consistent across different plaques of the same phage and differs between phages (**Figure 2**b). The initial plaque expansion rate correlates with the final plaque size (a), but this correlation is noisy (Spearman’s ρ=0.84, p<0.001, b) as the expansion start and final expansion rate also contribute to the variation in the final size (resp. Spearman’s ρ=0.26, p<0.001 and ρ=0.4, p<0.001, b). Therefore, the expansion dynamics offer insights into phenotypic diversity beyond end-point measurements.

We next wondered how phenotyping from plaque expansion compares to phenotyping in liquid assays. In liquid medium assays, bacterial growth is typically initially unperturbed by the presence of the phage. However, as the phage replicates, the bacterial population collapses at a time-point that is phage-dependent. We compared the time-point of collapse in a liquid LB-medium with the plaque expansion dynamics in a semi-solid LB-agar medium across the entire BASEL phage collection. We find that there is no correlation between the initial plaque expansion rate and the timepoint of collapse in liquid (Spearman’s ρ=0.07, p=0.25, b). Furthermore, there is only a weak correlation with the activity in liquid and the start of plaque expansion or the final expansion rate (resp. Spearman’s ρ=0.36, p<0.001 and ρ=0.31, p<0.001, b). Therefore, we conclude that plaque expansion dynamics and phage activity in liquid are governed by related but different processes.

We next wondered whether there were any notable trends in differences in plaque expansion dynamics between phages. We find that the initial expansion rate significantly decreases with the genome size (Spearman’s ρ=-0.7, p<0.001, b,c). This could be caused by smaller genomes requiring smaller capsids, and therefore a higher diffusion constant. Alternatively, this correlation could be caused by a faster replication rate of phages with a smaller genome. However, there is a confounding factor in that genome size correlates with the phage family (c). Indeed, the initial expansion rate not only correlates with genome size, it also significantly varies between phage families (d). When correcting for the phage family, no correlations can be found between the genome size and plaque expansion dynamics (Spearman’s ρ=-0.07, p=0.058, **Supplementary Figure 2**), making it impossible to disentangle the effects of genome size and phage family from our results.

As the phage family correlates strongly with the plaque expansion dynamics (c), we next wondered how accurately we can classify phages according to families solely from the plaque expansion dynamics. Therefore, we trained a random forest model to predict phage family from the five plaque expansion parameters. We find that using this approach we can accurately predict the phage family in 72.1% of the cases (**Figure 4**), without using any sequencing data. By contrast, using only end-point plaque sizes, we can only correctly assign the phage family in 40.4% of the cases, which is below the benchmark of naively guessing the largest phage family each time (49.1%, **Figure 4**). The costs of sequencing are low enough that this form of classification need not be replaced, however, the difference in classification power between end-point plaque size and the full plaque dynamics do highlight the power of our approach.

**Figure 4.**
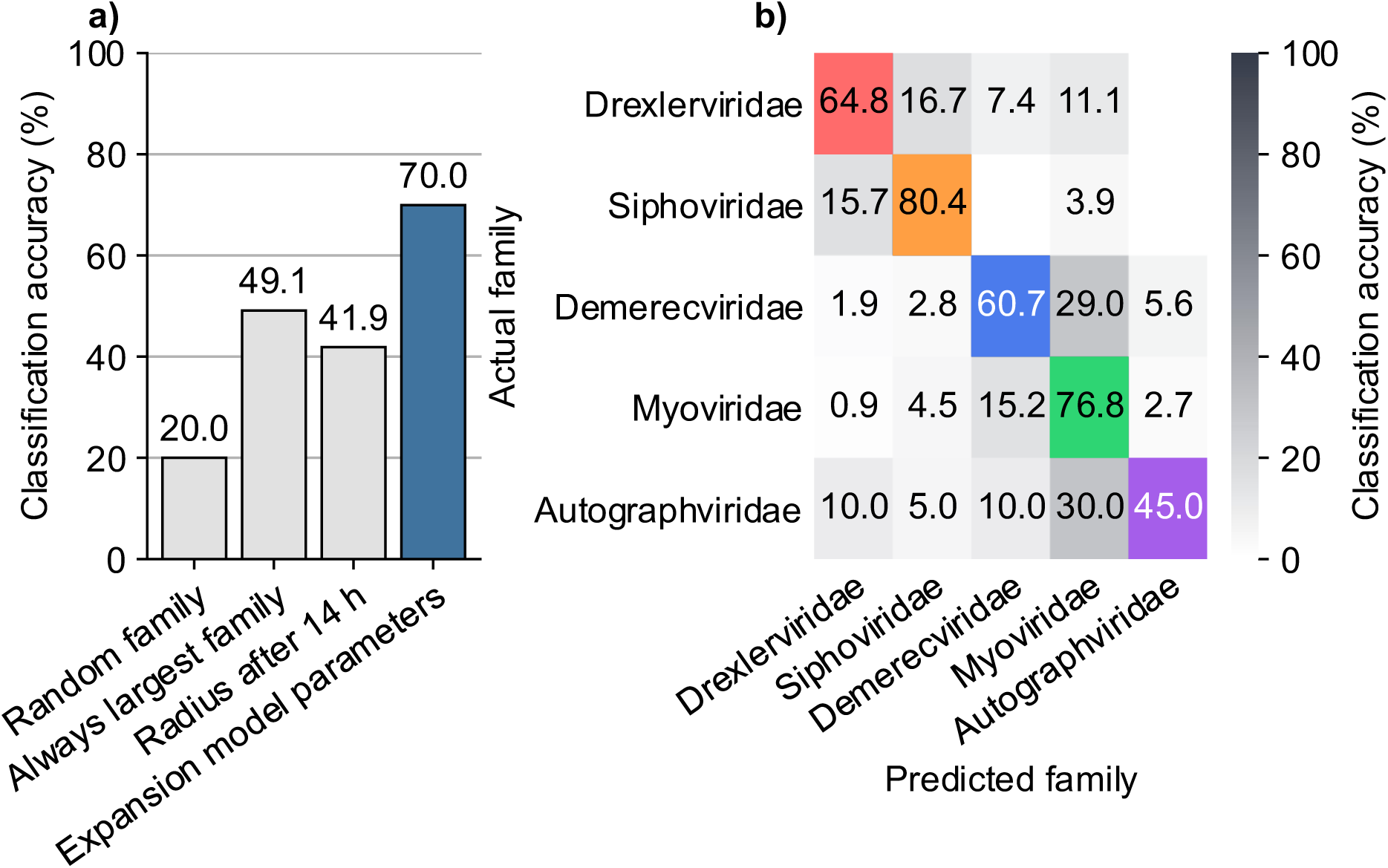
Phage family classification by plaque expansion dynamics. **a)** A random forest classifier was trained to assign phage families on basis of the plaque expansion dynamics and compared to a benchmark of randomly assigning a family or always assigning the largest family (myoviridae) to each phage. Whereas classification based on end-point plaque size alone performs below the benchmark of naively guessing the largest phage family each time, taking plaque dynamics into account correctly assigns the family to each phage in most cases. **b)** Detailed breakdown of classifications based on the full plaque dynamics. The colored diagonal shows the fraction of correct classifications per family.

## Discussion

We have used a consumer flatbed scanner combined with computer vision to track phage plaque dynamics. We have applied this assay to a diverse set of phages infecting the same host to find phage determinants of the plaque expansion dynamics. We found that the plaque expansion rate strongly correlates with both the phage family and the genome size, and that plaque dynamics are superior in classifying phages compared to end-point measurements. Therefore, we conclude that tracking plaque dynamics provides more detailed insights into phage phenotypes than end-point plaque size measurements.

Our open-source software package is useful beyond the scope of this work. Commercial flatbed scanners are widely used for scientific imaging, with applications including phage plaques^34^, bacterial colonies^46^ and applications outside of microbiology^47^. Communication typically relies on the TWAIN interface, which is a standardized software protocol to communicate with scanners and digital cameras. Our open-source software includes a connector to the standardized laboratory automation interface SiLA2^48^, enabling for ready-to-go integration of TWAIN-supported images in diverse laboratory automation pipelines^49,50^.

Up to now, assays measuring the bacterial killing kinetics of phages have been exclusively performed in liquid media. We find that the dynamics on agar correlate with but are different from the dynamics in liquid. We hypothesize that these differences are partly due to differences in the physiological state of bacteria in top agar and in liquid and partly because bacterial populations reach higher densities in top agar than in liquid. For therapeutic phage applications, it is unclear which better mimics the eradication dynamics of an infection. The Israeli phage therapy centers use both plaque assays and liquid media in selecting their phages^51^, but exclusively rely on liquid assays for kinetic information. Plaque expansion dynamics might offer a better proxy than liquid assays for bactericidal activity against biofilms due to their more similar cell densities and polymeric embedding. To quantitatively assess which assays best predict therapeutic success, we require more extensive retrospective clinical studies^52^ in combination with detailed in vitro characterization of the applied phages. Furthermore, preclinical models can help in causatively linking in vitro properties to therapeutic success^53^.

Beyond therapeutic applications, we also foresee several scientific applications for our method to track plaque expansion dynamics. There is a vast discrepancy in the diversity of phages we observe by sequencing and diversity of phages we culture and study in the lab. When sequencing a collection of isolated phages, one often finds the same or related phages^54^, thereby limiting the ultimate diversity of any phage bank. Recently, a similar problem for bacterial isolation was solved by using machine learning and automation to pick bacterial colonies with diverse morphologies to maximize the genetic diversity of bacterial isolates and enabled targeted picking of specific genera^55^. As we find a strong correlation between phage family and plaque expansion dynamics, we envision a similar approach can be used for phages. Therefore, our assay can be used as part of a broader pipeline to optimize phage discovery^54,56^.

Due to our limited understanding of phage biology, most phage genomes typically consist of genes with unknown functions. An especially promising direction in discovering functions at scale is by using explainable artificial intelligence (XAI) approaches on large data sets to link phenotypes and genotypes^57^. This XAI approach has been recently applied to phages^58^, but the success of these models relies heavily on the available data, which for phages is fully dominated by relatively crude, binary phenotypes of infecting/non-infecting^6–10^. Most phage genes do not play an essential role in infectivity^59^, but likely still affect the rate of bacterial killing. We believe that by scaling up our assay and applying it to many combinations of different phages and bacteria the power of XAI can be expanded to unravelling the function of diverse non-essential genes.

## Declaration of generative AI and AI-assisted technologies in the manuscript preparation process

During the preparation of this work the authors used Gemini in order to optimize the writing for accessibility. After using this tool, the authors reviewed and edited the content as needed and take full responsibility for the content of the published article.

## Funding

This research did not receive any specific grant from funding agencies in the public, commercial, or not-for-profit sectors.

## Author contributions

Conceptualization: JC, YM, Data curation: JC, formal analysis: JC, Investigation: JC, BAS, Methodology: JC, BAS, YM, software: JC, supervision: BAS, YM, Visualization: JC, Writing – original draft: JC, YM, writing – review and editing: JC, BAS, YM

## Acknowledgements

We thank the Molecular Microbiology section of the Vrije Universiteit Amsterdam for their valuable input throughout the project.

## Supplementary Figures

**Supplementary Figure 1.**
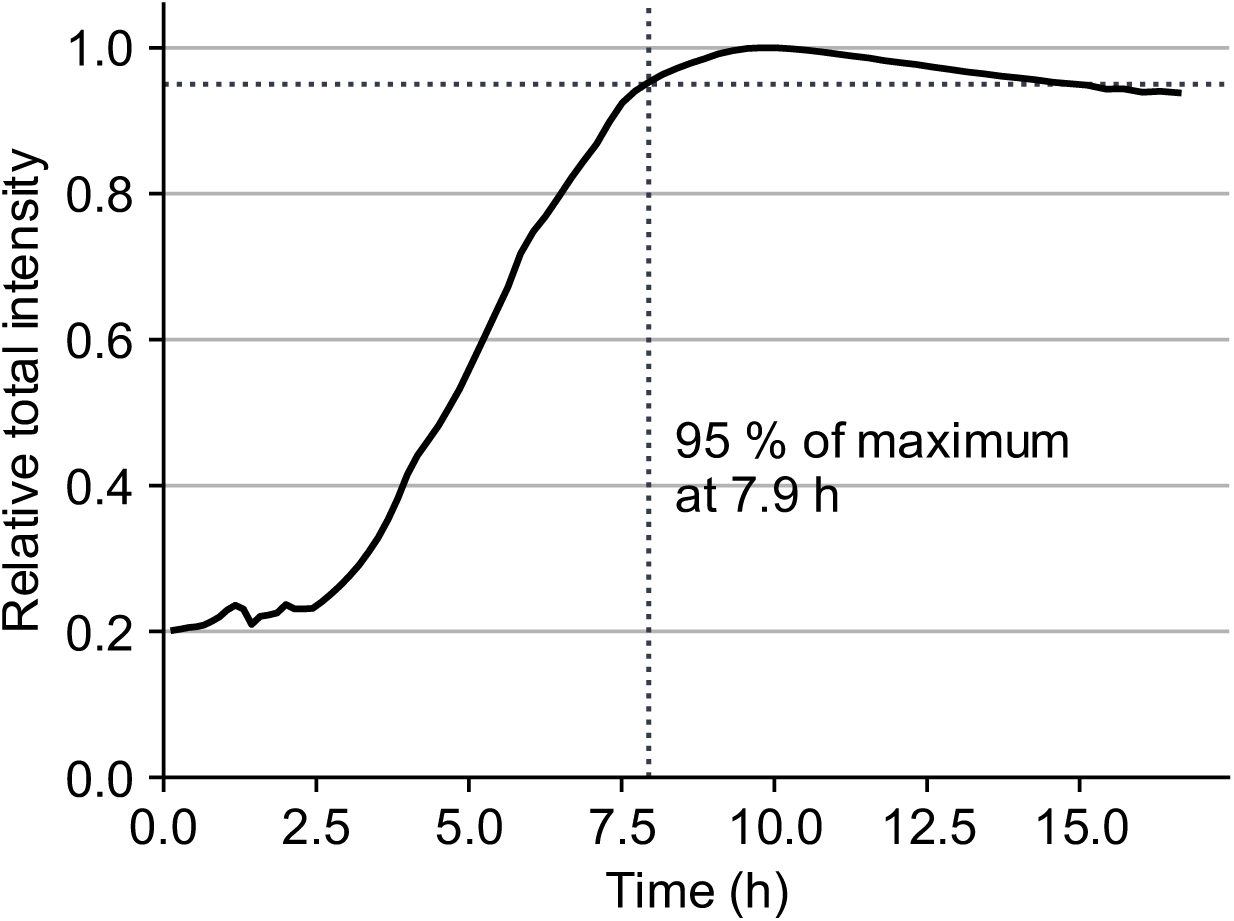
Bacterial growth saturates after 8 hours. Bacterial growth was quantified by integrating pixel intensity over a plaque-free region, normalized to the maximum intensity recorded across the time-lapse. Plates were pre-incubated to minimize the lag phase. Signal scattering from the bacterial lawn exceeded background levels after 2.5 hours and increased approximately linearly until saturating after roughly 8 hours.

**Supplementary Figure 2.**
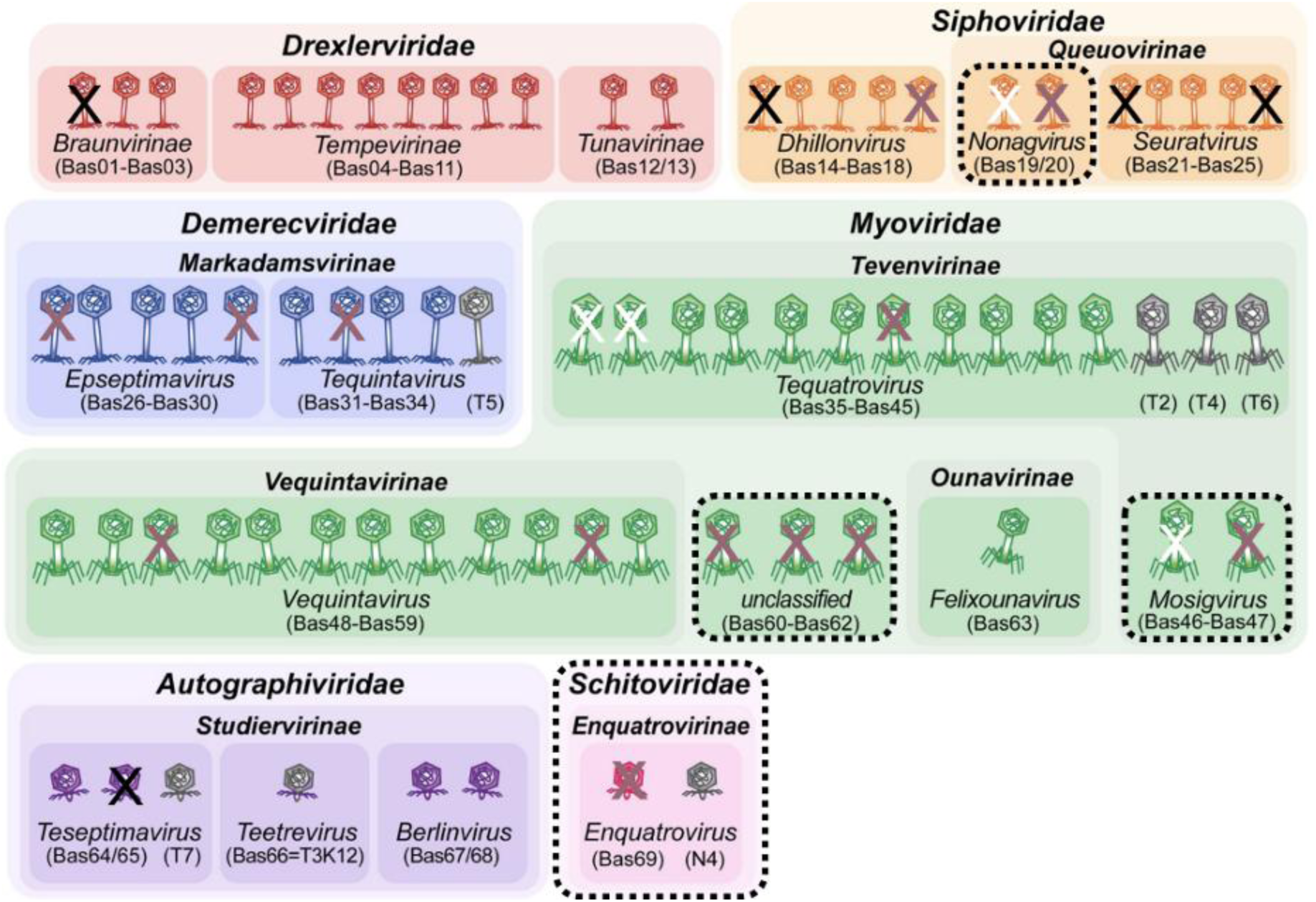
Visual representation of the phages present in the BASEL library overlaid with marks for uncaptured phages. Adapted from Ref. ^31^. Phages crossed out in dark pink mark loss due to size, in black loss due to abnormality and white loss due to absence from stock. The families and subfamilies fully uncaptured are marked with black dashed lines. Grayed out phages are not part of the library.

**Supplementary Figure 3.**
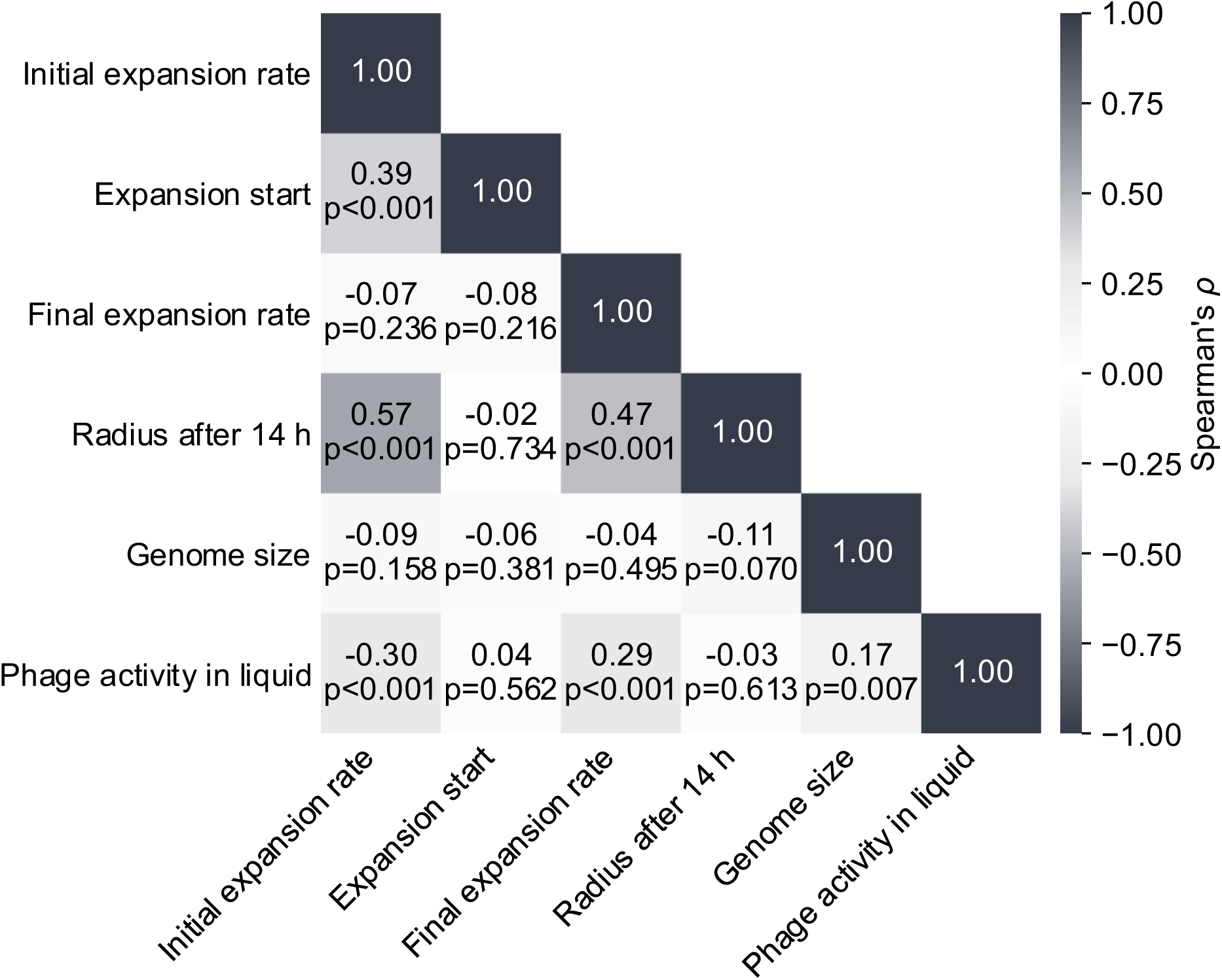
Partial Spearman correlation matrix corrected for phage family. Phage family and genome size are strongly correlated. When correcting for the phage family, the genome size no longer correlates with either the initial expansion rate or the expansion start. Therefore, it is not possible to decouple genome size effects from phage family effects.

## Notes

https://zenodo.org/records/23216050

### Competing Interest Statement

The authors have declared no competing interest.

## References

1. Dion, M. B., Oechslin, F. & Moineau, S. Phage diversity, genomics and phylogeny. Nat. Rev. Microbiol. 18, 125–138 (2020).

2. Wei, C. & Chen, Z. Comprehensive analysis of phage genomes from diverse environments reveals their diversity, potential applications, and interactions with hosts and other phages. Front. Microbiol. 16, (2025).

3. Blackwell, G. A. et al. Exploring bacterial diversity via a curated and searchable snapshot of archived DNA sequences. PLOS Biol. 19, e3001421 (2021).

4. Leprince, A., Somerville, V., Addablah, A. A., Morency, C. & Moineau, S. Phage host range: determinants, dynamics and applications. Nat. Rev. Microbiol. 1–21 (2026) doi:10.1038/s41579-026-01306-x.

5. Mulla, Y., Müller, J., Trimcev, D. & Bollenbach, T. Extreme diversity of phage amplification rates and phage–antibiotic interactions revealed by PHORCE. PLOS Biol. 23, e3003065 (2025).

6. Costa, A. R. et al. Accumulation of defense systems in phage-resistant strains of Pseudomonas aeruginosa. Sci. Adv. 10, eadj0341 (2024).

7. Rothschild-Rodriguez, D. et al. KlebPhaCol: a community-driven resource for *Klebsiella* research identified a novel phage family. Nucleic Acids Res. 53, gkaf1122 (2025).

8. Kauffman, K. M. et al. Resolving the structure of phage–bacteria interactions in the context of natural diversity. Nat. Commun. 13, 372 (2022).

9. Piel, D. et al. Phage–host coevolution in natural populations. Nat. Microbiol. 7, 1075–1086 (2022).

10. Gaborieau, B. et al. Prediction of strain level phage–host interactions across the Escherichia genus using only genomic information. Nat. Microbiol. 9, 2847–2861 (2024).

11. Leung, C. Y. (Joey) & Weitz, J. S. Modeling the synergistic elimination of bacteria by phage and the innate immune system. J. Theor. Biol. 429, 241–252 (2017).

12. Van Der Steen, B. A., Gralka, M. & Mulla, Y. Rapid resistance evolution against phage cocktails. ISME J. 20, wrag067 (2026).

13. Glonti, T. & Pirnay, J.-P. In Vitro Techniques and Measurements of Phage Characteristics That Are Important for Phage Therapy Success. Viruses 14, 1490 (2022).

14. Konopacki, M., Grygorcewicz, B., Dołęgowska, B., Kordas, M. & Rakoczy, R. PhageScore: A simple method for comparative evaluation of bacteriophages lytic activity. Biochem. Eng. J. 161, 107652 (2020).

15. Storms, Z. J., Teel, M. R., Mercurio, K. & Sauvageau, D. The Virulence Index: A Metric for Quantitative Analysis of Phage Virulence. PHAGE 1, 27–36 (2020).

16. Kolenda, C., Jourdan, J., Roussel-Gaillard, T., Medina, M. & Laurent, F. Phage susceptibility testing methods or ‘phagograms’: where do we stand and where should we go? J. Antimicrob. Chemother. dkae325 (2024) doi:10.1093/jac/dkae325.

17. Xie, Y. et al. Differential Bacteriophage Efficacy in Controlling Salmonella in Cattle Hide and Soil Models. Front. Microbiol. 12, (2021).

18. Kutter, E. Phage Host Range and Efficiency of Plating. in Bacteriophages: Methods and Protocols, Volume 1: Isolation, Characterization, and Interactions (eds Clokie, M. R. J. & Kropinski, A. M.) 141–149 (Humana Press, Totowa, NJ, 2009). doi:10.1007/978-1-60327-164-6_14.

19. Burch, C. L. & Chao, L. Epistasis and Its Relationship to Canalization in the RNA Virus φ6. Genetics 167, 559–567 (2004).

20. Roychoudhury, P., Shrestha, N., Wiss, V. R. & Krone, S. M. Fitness benefits of low infectivity in a spatially structured population of bacteriophages. Proc. R. Soc. B Biol. Sci. 281, 20132563 (2014).

21. Bulssico, J., Papukashvili, I., Espinosa, L., Gandon, S. & Ansaldi, M. Phage-Antibiotic Synergy: Cell Filamentation Is a Key Driver of Successful Phage Predation. http://biorxiv.org/lookup/doi/10.1101/2022.11.28.518157 (2022) doi:10.1101/2022.11.28.518157.

22. Nabergoj, D., Modic, P. & Podgornik, A. Effect of bacterial growth rate on bacteriophage population growth rate. MicrobiologyOpen 7, e00558 (2018).

23. Gallet, R., Kannoly, S. & Wang, I.-N. Effects of bacteriophage traits on plaque formation. BMC Microbiol. 11, 181 (2011).

24. Ortega-Cejas, V., Fort, J., Méndez, V. & Campos, D. Approximate solution to the speed of spreading viruses. Phys. Rev. E Stat. Nonlin. Soft Matter Phys. 69, 031909 (2004).

25. Fort, J. & Méndez, V. Time-Delayed Spread of Viruses in Growing Plaques. Phys. Rev. Lett. 89, 178101 (2002).

26. Abedon, S. T. & Yin, J. Bacteriophage Plaques: Theory and Analysis. Methods Mol. Biol. 10.1007/978-1-60327-164-6_17 (2009) doi:10.1007/978-1-60327-164-6_17.

27. Hunter, M., Krishnan, N., Liu, T., Möbius, W. & Fusco, D. Virus-Host Interactions Shape Viral Dispersal Giving Rise to Distinct Classes of Traveling Waves in Spatial Expansions. Phys. Rev. X 11, 021066 (2021).

28. Yin, J. Evolution of bacteriophage T7 in a growing plaque. J. Bacteriol. 175, 1272–1277 (1993).

29. Lee, Y. & Yin, J. Imaging the propagation of viruses. Biotechnol. Bioeng. 52, 438–442 (1996).

30. Yin, J. A quantifiable phenotype of viral propagation. Biochem. Biophys. Res. Commun. 174, 1009–1014 (1991).

31. Maffei, E. et al. Systematic exploration of Escherichia coli phage–host interactions with the BASEL phage collection. PLOS Biol. 19, e3001424 (2021).

32. Alam, H. & Fusco, D. Resource availability and dimensionality result in ecology-dependent selection in bacteriophage spatial expansions. 2026.02.23.707387 Preprint at 10.64898/2026.02.23.707387 (2026).

33. Sinha, S., Grewal, R. K. & Roy, S. Modeling Bacteria–Phage Interactions and Its Implications for Phage Therapy. in Advances in Applied Microbiology vol. 103 103–141 (Elsevier, 2018).

34. Payne, P., Geyrhofer, L., Barton, N. H. & Bollback, J. P. CRISPR-based herd immunity can limit phage epidemics in bacterial populations. eLife 7, e32035 (2018).

35. Möbius, W., Murray, A. W. & Nelson, D. R. How Obstacles Perturb Population Fronts and Alter Their Genetic Structure. PLoS Comput. Biol. 11, e1004615 (2015).

36. Brewster, J. D. A simple micro-growth assay for enumerating bacteria. J. Microbiol. Methods 53, 77–86 (2003).

37. TWAIN Working Group. https://twain.org/?mid=17Cid=43Ctid=15#/user/home.

38. UniteLabs / Connector Development Kit / Python CDK · GitLab. GitLab https://gitlab.com/unitelabs/cdk/python-cdk (2026).

39. Juchli, D. SiLA 2: The Next Generation Lab Automation Standard. Adv. Biochem. Eng. Biotechnol. 182, 147–174 (2022).

40. SiLA Rapid Integration. https://sila-standard.com/ (2017).

41. Jonathan Costa / Plaque Dynamics · GitLab. GitLab https://gitlab.com/costa-jonathan/plaque-dynamics (2026).

42. Hoyer, S. & Hamman, J. xarray: N-D labeled Arrays and Datasets in Python. J. Open Res. Softw. 5, (2017).

43. GitHub - zarr-developers/zarr-python: An implementation of chunked, compressed, N-dimensional arrays for Python. · GitHub. https://github.com/zarr-developers/zarr-python.

44. astral-sh/uv. Astral (2026).

45. Walt, S. van der et al. scikit-image: Image processing in Python. PeerJ 2, e453 (2014).

46. Levin-Reisman, I. et al. Automated imaging with ScanLag reveals previously undetectable bacterial growth phenotypes. Nat. Methods 7, 737–739 (2010).

47. Mohamed, A. et al. An evaluation of inexpensive methods for root image acquisition when using rhizotrons. Plant Methods 13, 11 (2017).

48. Bromig, L. et al. The SiLA 2 Manager for rapid device integration and workflow automation. SoftwareX 17, 100991 (2022).

49. Sim, M. et al. ChemOS 2.0: An orchestration architecture for chemical self-driving laboratories. Matter 7, 2959–2977 (2024).

50. Courtney, P. et al. An out-of-hours cell culture autopilot proof of concept based on a modular architecture using the SiLA2 open standard. SLAS Technol. 32, 100279 (2025).

51. Yerushalmy, O. et al. Towards Standardization of Phage Susceptibility Testing: The Israeli Phage Therapy Center “Clinical Phage Microbiology”—A Pipeline Proposal. Clin. Infect. Dis. 77, S337–S351 (2023).

52. Pirnay, J.-P. et al. Personalized bacteriophage therapy outcomes for 100 consecutive cases: a multicentre, multinational, retrospective observational study. Nat. Microbiol. 9, 1434–1453 (2024).

53. Lindberg, H. M., McKean, K. A. & Wang, I.-N. Phage fitness may help predict phage therapy efficacy. Bacteriophage 4, e964081 (2014).

54. Olsen, N. S., Hendriksen, N. B., Hansen, L. H. & Kot, W. A New High-Throughput Screening Method for Phages: Enabling Crude Isolation and Fast Identification of Diverse Phages with Therapeutic Potential. Phage 1, 137–148 (2020).

55. Huang, Y. et al. High-throughput microbial culturomics using automation and machine learning. Nat. Biotechnol. 41, 1424–1433 (2023).

56. Phage DisCo: targeted discovery of bacteriophages by co-culture | mSystems. https://journals.asm.org/doi/10.1128/msystems.01644-24.

57. Ma, J. et al. Using deep learning to model the hierarchical structure and function of a cell. Nat. Methods 15, 290–298 (2018).

58. Noonan, A. J. C. et al. Phylogeny-agnostic strain-level prediction of phage-host interactions from genomes. 2025.11.15.688630 Preprint at 10.1101/2025.11.15.688630 (2026).

59. Chen, J. et al. Systematic, high-throughput characterization of bacteriophage gene essentiality on diverse hosts. Cell Host Microbe 33, 1363–1380.e11 (2025).

